# Disentangling the effects of selection and loss bias on gene dynamics

**DOI:** 10.1101/139725

**Authors:** Jaime Iranzo, José A. Cuesta, Susanna Manrubia, Mikhail I. Katsnelson, Eugene V. Koonin

**Affiliations:** National Center for Biotechnology Information, National Library of Medicine, National Institutesof 6 Health, Bethesda, MD 20894, USA; Grupo Interdisciplinar de Sistemas Complejos (GISC); Departamento de Matemáticas, Universidad Carlos III de Madrid, Spain; Institute for Biocomputation and Physics of Complex Systems, Zaragoza, Spain; UC3M-BS Institute of Financial Big Data (IFiBiD); Grupo Interdisciplinar de Sistemas Complejos (GISC); National Biotechnology Centre (CSIC), Madrd,Spain; Institute for Molecules and Materials, Radboud University, Nijmegen, 6525AJ, Netherlands.

## Abstract

We combine mathematical modelling of genome evolution with comparative analysis of prokaryotic genomes to estimate the relative contributions of selection and intrinsic loss bias to the evolution of different functional classes of genes and mobile genetic elements (MGE). An exact solution for the dynamics of gene family size was obtained under a linear duplication-transfer-loss model with selection. With the exception of genes involved in information processing, particularly translation, which are maintained by strong selection, the average selection coefficient for most non-parasitic genes is low albeit positive, compatible with the observed positive correlation between genome size and effective population size. Free-living microbes evolve under stronger selection for gene retention than parasites. Different classes of MGE show a broad range of fitness effects, from the nearly neutral transposons to prophages, which are actively eliminated by selection. Genes involved in anti-parasite defense, on average, incur a fitness cost to the host that is at least as high as the cost of plasmids. This cost is probably due to the adverse effects of autoimmunity and curtailment of horizontal gene transfer caused by the defense systems and selfish behavior of some of these systems, such as toxin-antitoxin and restriction-modification modules. Transposons follow a biphasic dynamics, with bursts of gene proliferation followed by decay in the copy number that is quantitatively captured by the model. The horizontal gene transfer to loss ratio, but not the duplication to loss ratio, correlates with genome size, potentially explaining the increased abundance of neutral and costly elements in larger genomes.

**SIGNIFICANCE:** Evolution of microbes is dominated by horizontal gene transfer and the incessant host-parasite arms race that promotes the evolution of diverse anti-parasite defense systems. The evolutionary factors governing these processes are complex and difficult to disentangle but the rapidly growing genome databases provide ample material for testing evolutionary models. Rigorous mathematical modeling of evolutionary processes, combined with computer simulation and comparative genomics, allowed us to elucidate the evolutionary regimes of different classes of microbial genes. Only genes involved in key informational and metabolic pathways are subject to strong selection whereas most of the others are effectively neutral or even burdensome. Mobile genetic elements and defense systems are costly, supporting the understanding that their evolution is governed by the same factors.

## Introduction

In the wake of the genomic revolution, quantitative understanding of the roles that ecological and genetic factors play in determining the size, composition and architecture of genomes has become a central goal in biology (1-3). The vast number of prokaryotic genomes sequenced to date reveals a great diversity of sizes, which range from about 110kb and 140 protein coding genes in the smallest intracellular symbionts (4) to almost 15Mb and more than 10,000 genes in the largest myxobacteria (5). Beyond a core of approximately 100 nearly universal genes, the gene complements of bacteria and archaea are highly heterogeneous (6-8). Remarkably, 10-20% of the genes in most microbial genomes are ORFans, that is, genes that have no detectable homologs in other species and are replaced at extremely high rates in the course of microbial evolution (9, 10). Furthermore, all but the most reduced genomes host multiple and diverse parasitic genetic elements, such as transposons and prophages that collectively comprise the so-called microbial mobilome (11).

The evolution of microbial genomes is generally interpreted in terms of the interplay between three factors: i) gene gain, via horizontal gene transfer (HGT) and gene duplication, ii) gene loss, via deletion, and iii) natural selection that affects the fixation and maintenance of genes (8, 12). The intrinsic bias towards DNA deletion (and hence gene loss) that characterizes mutational processes in prokaryotes (as well as eukaryotes) results in non-adaptive genome reduction (13), whereas selection contributes to maintaining slightly beneficial genes (14). In agreement with this model, the strength of purifying selection, as measured by the ratio of non-synonymous to synonymous variation, positively correlates with the genome size (15, 16). However, when it comes to interpreting the genome composition, the picture is complicated by the fact that selection can also lead to adaptive genome reduction by removing pseudogenes (17), costly genetic parasites, and accessory genes, which are dispensable under stable environmental conditions (18, 19). Conversely, the increased propensity of some gene families to be horizontally transferred might suffice to ensure their persistence beyond the effects of selection and intrinsic loss bias (20). Rather than being minor deviations from a general trend, non-uniform levels of selection and horizontal gene transfer affecting different families and classes of genes appear to be essential to explain the abundance distributions and evolutionary persistence times of genes (10, 12). Accordingly, a quantitative assessment of the fitness costs and benefits for different classes of genes is essential to attain an adequate understanding of the evolutionary forces that shape genomes.

The magnitude and even the sign with which the presence (or absence) of a gene contributes to the fitness of an organism are not constant in time. For example, the metabolic cost incurred by the replication, transcription and translation of a gene strongly depends on the cell growth rate and the gene expression level (21). A recent study on the effects of different types of mutations in *Salmonella enterica* has shown that up to 25% of large deletions could result in a fitness increase although the benefit of losing a particular gene critically depends on the environment (19). These findings emphasize the importance of averaging across multiple environmental conditions when it comes to estimating the fitness contribution of a gene. For the purpose of evolutionary analyses, a meaningful proxy for such an average can be obtained by inferring selection coefficients directly from the gene family abundances observed in large collections of genomes. The main difficulty in this case is disentangling the effects of selection from the effects of intrinsic loss bias, which normally requires a priori knowledge of the effective population size or the gene gain and loss rates (14, 22, 23).

Here we combine mathematical modelling, comparative genomics, and data compiled from mutation accumulation experiments to infer the characteristic contributions of selection and intrinsic DNA loss for different gene categories. To disentangle selection and loss bias, we first obtained an exact, time-dependent solution of the linear duplication-transfer-loss model with selection that governs the dynamics of gene copy numbers in a population of genomes (24-28). When applied to a large genomic data set, the model provides maximum likelihood estimates of the ‘neutral equivalent’ (‘effective’) loss bias, a composite parameter that amalgamates the effects of intrinsic loss bias (the loss bias prior to the action of selection) and selection. The selection coefficient can be extracted from the effective loss bias as long as the rate of gene loss is known, for which we used estimates from mutation accumulation experiments.

Our results show that, with the exception of genes involved in core informational processes, most gene families are neutral or only slightly beneficial in the long term. Among the genetic elements that are typically considered parasitic, prophages show the highest fitness cost, followed by conjugative plasmids and transposons, which are only weakly deleterious in the long term. Notably, genes involved in anti-parasite defense do not seem to provide long-term benefits on average, but rather are slightly deleterious, almost to the same extent as transposons. We complete our analysis with an evaluation of the causes that make transposon dynamics qualitatively different from that of other gene classes and explore the effect of genome size on the rates of HGT, gene duplication and gene loss.

## Results

### Duplication-transfer-loss model of gene family evolution

To describe the dynamics of a gene family size (gene copy number) in a population of genomes, we employed a linear duplication-transfer-loss model with selection. Within a genome, the gene copy number can increase via duplication of the extant copies, which occurs at rate *d* per copy, or through the arrival of a new copy via HGT, at rate *h* independent on the copy number. Likewise, gene loss at rate *l* per copy leads to a decrease in the copy number. Duplication, HGT and gene loss define a classical birth-death-transfer model at the genome level (24-27, 29). Selection is introduced through a contribution *s* to the fitness of a genome (*s* is positive for beneficial genes and negative for costly genes), which is multiplied by the gene copy number *k*. Specifically, we assume that fitness is additive, there is no epistasis, and the fitness contributions of all genes from the same family are the same. At the cell population level, the number of genomes carrying *k* copies, *n*_*k*_, obeys the following system of differential equations:

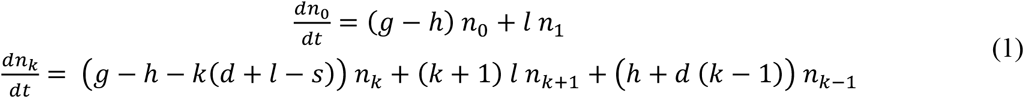

The basal growth rate *g* was included for completeness although it does not affect the copy number distribution. Moreover, the entire system can be restated in terms of the ratios of each of the parameters to the loss rate (see SI text for more details). The linear duplication-transfer-loss model with selection can be exactly solved for arbitrary initial conditions by formulating eq. (1) as a first-order partial differential equation for the generating function and applying the method of characteristics (SI text)(30, 31). The result is the copy number distribution, i.e. the fraction *p*_*k*_ of hosts with an arbitrary number of copies *k* at any time. In the case of a population where the gene family is initially absent, we obtain

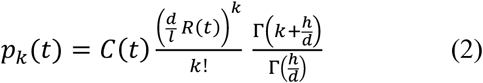

with 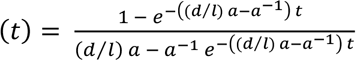, and 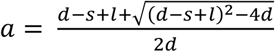. In these expressions, time is measured in units of loss events and *C*(*t*) is a normalization factor that ensures that the sum of *p*_*k*_ over all *k* is equal to 1. A notable property of this solution is that, as the system approaches the stationary state,selection, duplication and loss merge into the composite parameter *a* which, in the absence of selection, coincides with the inverse of the duplication/loss ratio (see SI text for more details). Therefore, we refer to *a*^−1^ as the ‘neutral equivalent’ (henceforth ‘effective’) duplication/loss ratio (*d*/*l*_*e*_). It is also possible to define the effective HGT/loss ratio 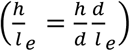 such that gene families with the same effective ratios have the same stationary distributions. The fitness contribution of a gene (i.e. selection to loss ratio) can be expressed in terms of the gene’s effective duplication/loss ratio and the actual (intrinsic)duplication/loss ratio as

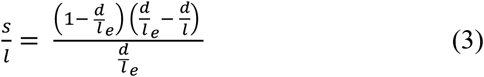

### Duplication, loss and selection in different functional categories of genes

We used the COUNT method to estimate the effective duplication/loss ratio (*d*/*l*_*e*_) associated to different gene families (defined as Clusters of Orthologous Groups, or COGs) in 35 sets of closely related genomes (Alignable Tight Genomic Clusters, or ATGCs), which jointly encompass 678 bacterial and archaeal genomes (32, 33). As shown in the preceding section, the effective duplication/loss ratio (*d*/*l*_e_) is a composite parameter that results from selection on gene copy number affecting the fixation of gene duplications and gene losses. For a neutral gene family, the effective duplication/loss ratio is simply the same as the ratio between the rates of gene duplication and gene loss. Because selection prevents the loss of beneficial genes, the effective duplication/loss ratios associated with beneficial genes are greater than their intrinsic duplication/loss ratios, whereas the opposite holds for genes (e.g. parasitic elements) that are costly to the host and tend to be eliminated by selection. Technically, the duplication term includes not only *bona fide* duplications but any process that causes an increase in copy number that is proportional to the preexisting copy number. Thus, HGT can also contribute to the duplication term in clonal populations, where the copy numbers of donors and recipients are highly correlated. Fig. 1A shows the effective duplication/loss ratios for gene families that belong to different functional categories (as defined under the COG classification (34)), as well as genes of transposons, conjugative plasmids and prophages. For the majority of the gene families, the effective duplication/loss ratios are below 1, which is compatible with the pervasive bias towards gene loss combined with (near) neutrality of numerous genes. In agreement with the notion that selection affects the effective duplication/loss ratios, their values decrease from the essential functional categories, such as translation and nucleotide metabolism, to the non-essential and parasitic gene classes. The apparent bimodality of the distributions for some functional categories (Fig. 1A) is likely due to their biological heterogeneity. For example, category N (secretion and motility), sharply splits into two major groups of gene families: (i) components of the flagellum and (ii)proteins involved in cellulose production and glycosyltransferases, with high *d*/*l*_*e*_ values for the former and much lower values for the latter (SI Table S1).

**Figure 1:**
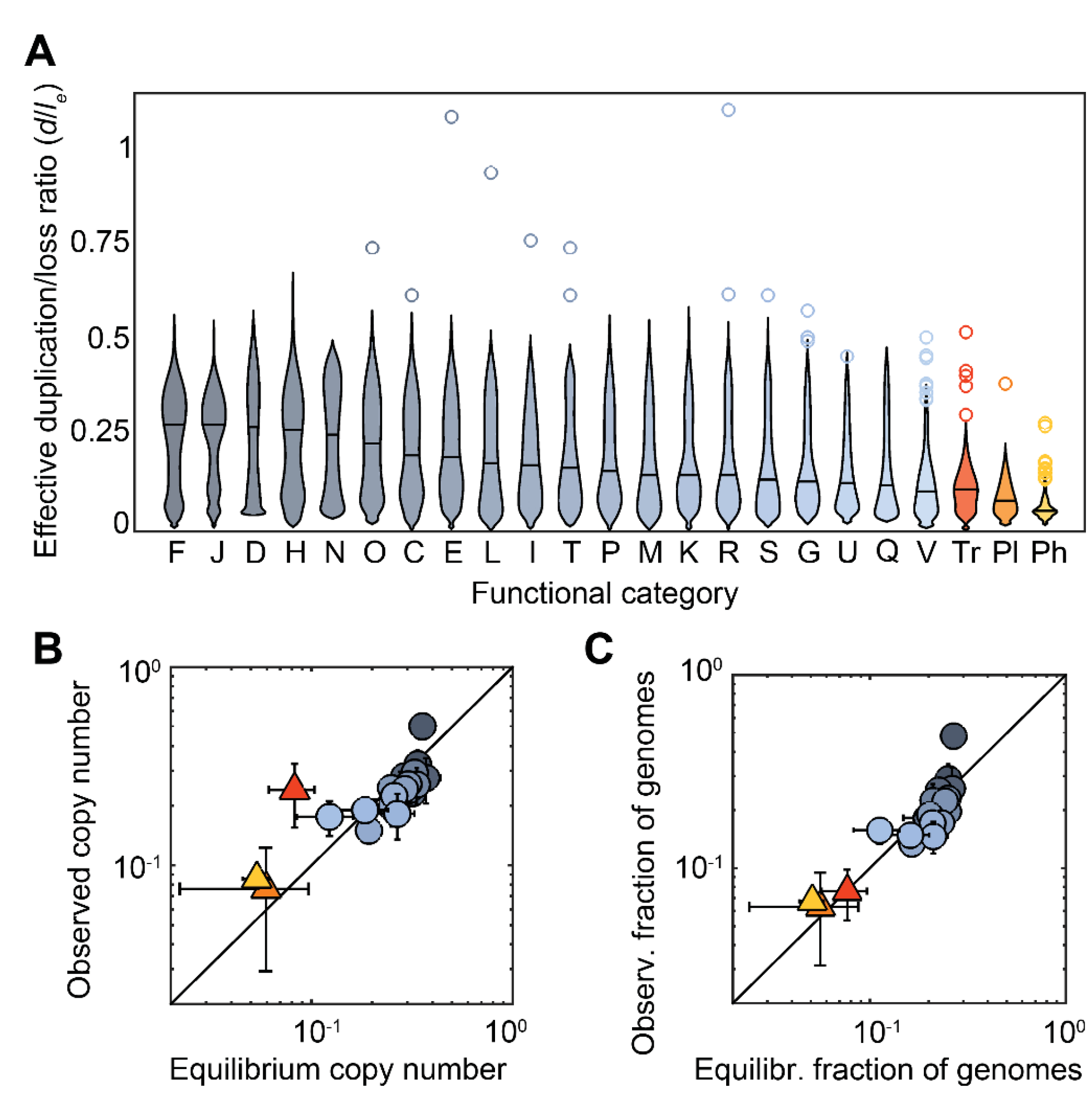
Effective loss bias and mean abundances of gene families from different functional categories. A: Distribution of the effective duplication/loss ratio *d*/*l*_*e*_. Black horizontal lines indicate the median of each category. Outliers are represented as circles. Designations of the functional categories (modified from (8)): C, energy production and conversion; D, cell division; E, amino acid metabolism and transport; F, nucleotide metabolism and transport; G, carbohydrate metabolism and transport; H, coenzyme metabolism; I, lipid metabolism; J, translation; K, transcription; L, replication and repair; M, membrane and cell wall structure and biogenesis; N, secretion and motility; O, post-translational modification, protein turnover and chaperone functions; P, inorganic ion transport and metabolism; Q, biosynthesis, transport and catabolism of secondary metabolites; R, general functional prediction only (typically, prediction of biochemical activity); S, function unknown; T, signal transduction; U, intracellular trafficking and secretion; V, defense mechanisms; Tr, transposon; Pl, conjugative plasmid; Ph, prophage or phage-related. Two extreme outliers, one from the transposons (transposase IS1595, *d*/*l*_*e*_ = 1.4) and one from category V (multidrug efflux pump subunit AcrB, *d*/*l*_*e*_ = 1.6) are not represented. B: Comparison of the global (observed) mean copy number per family and the equilibrium copy number predicted by the model. Data points correspond to medians across functional categories (colors as in A, triangles are used to highlight genetic parasites). Error bars represent the 95% confidence interval for the median. The solid line corresponds to a perfect match between predictions and observations. The Spearman’s correlation coefficients including and excluding parasites are *ρ* = 0.80 and 0.81, respectively (*p*<10^−4^). C: Fraction of genomes in which a family is present, compared with the expected fraction at equilibrium (Spearman’s *ρ* = 0.87 and 0.80, including and excluding parasites, *p* > 10^−4^). Data points and error bars as in B.

The average fitness contribution of a gene can be inferred from its effective duplication/loss ratio provided that the intrinsic duplication/loss ratio is known (see preceding section). To estimate the intrinsic duplication/loss ratio (*d*/*l*), we employed two independent approaches. The first approach was based on the assumption that a substantial fraction of genes from non-essential, but not parasitic, functional categories are effectively neutral. Considering that gene families in those categories are relatively well represented across taxa (we required them to be present in at least 3 different ATGCs) and are not regarded as part of the mobilome (11), we would expect that, if not neutral, they are slightly beneficial and provide an upper bound for the intrinsic duplication/loss ratio. After sorting non-parasitic functional categories by their effective duplication/loss ratios (Fig. 1A), category K (transcription) was selected as the last category whose members arguably exert a positive average fitness effect. The intrinsic duplication/loss ratio was then calculated as the median of the effective duplication/loss ratios among the pool of gene families involved in poorly understood functions (R, S), carbohydrate metabolism (G), secretion (U), secondary metabolism (Q), and defense (V). In the second approach, we identified genes that are represented by one or more copies in a single genome, while absent in all other genomes of the same ATGC. Such genes (henceforth ORFans (35, 36)) are likely of recent acquisition and can be assumed neutral, if not slightly deleterious. The maximum likelihood estimate of the duplication/loss ratio obtained for ORFans provides, therefore, a lower bound for the intrinsic duplication/loss ratio (see Methods and SI text). The ratios obtained with both approaches were 0.124 (95% CI 0.117-0.131) and 0.126 (95% CI 0.115-0.137), respectively. The two independent estimates are strikingly consistent with each other and robust to small changes in the methodology (SI text). Accordingly, we took the average *d*/*l* = 0.125 as the intrinsic duplication/loss ratio. This value quantifies the intrinsic bias towards gene loss once the effect of selection is removed.

Quantitative estimates of the ratio between the selection coefficient and the loss rate (*s*/*l*) for each functional category are readily obtained by applying eq. (3) to the effective duplication/loss ratios (Table 1). In the case of costly gene families, the ratio *s*/*l* quantifies the relative contributions of selection and loss in controlling the gene copy number. However, quantitative assessment of the selection coefficients from the *s*/*l* ratio requires knowledge of the intrinsic rates of gene loss in prokaryotic genomes. A compilation of published data from mutation accumulation experiments shows that disruption of gene coding regions due to small indels and/or large deletions occurs at rates between 5 × 10^−9^ and 4 ×10^−8^ per gene per generation (37-45), which yields the ranges for the selection coefficients listed in Table 1.Assuming the effective size of typical microbial populations to fall between 10^8^ and 10^9^ (21, 46, 47), the selection coefficients yielded by these estimates indicate evolution determined by positive fitness contribution (*N*_*e*_*s* ≫ 1) for information processing categories (translation and replication) as well as some metabolic categories (especially, nucleotide metabolism) and cellular functions (cell division,chaperones); an effectively neutral evolutionary regime for several categories including transcription; and evolution driven by negative fitness contribution (*N*_*e*_*s* ≪ −1) for defense genes and mobile genetic elements.

**Table 1:** Contributions of selection and the duplication/loss ratio to the evolution of different functional categories of genes and mobile elements The table shows the estimated values of the effective duplication/loss ratio (*d*/*l*_*e*_), selection to loss ratio(*s/l*) and fitness cost (selection coefficient) (*s*) for different functional categories of genes The *s*/*l* values were calculated assuming an intrinsic duplication/loss ratio *d*/*l* = 0.125. Loss rates equal to 5 × and4 × 1 per gene per generation were used to obtain the lower and upper estimates of *s*, respectively.

| | $d/l_e$ | $s/l$ | $s (\times 10^{-8})$ | |
| --- | --- | --- | --- | --- |
|  |  |  | Lower | upper |
| F - nucleotide metabolism and transport | 0.273 | 0.39 | 0.20 | 1.58 |
| J – translation | 0.273 | 0.39 | 0.20 | 1.58 |
| D - cell division | 0.266 | 0.39 | 0.19 | 1.56 |
| H - coenzyme metabolism | 0.260 | 0.38 | 0.19 | 1.54 |
| N - secretion and motility | 0.247 | 0.37 | 0.19 | 1.49 |
| O - post-translational modification, protein turnover and chaperone functions | 0.223 | 0.34 | 0.17 | 1.37 |
| C - energy production and conversion | 0.197 | 0.29 | 0.15 | 1.18 |
| E - amino acid metabolism and transport | 0.187 | 0.27 | 0.14 | 1.08 |
| L - replication and repair | 0.172 | 0.23 | 0.11 | 0.91 |
| I - lipid metabolism | 0.166 | 0.20 | 0.10 | 0.82 |
| T - signal transduction | 0.159 | 0.18 | 0.09 | 0.72 |
| P - inorganic ion transport and metabolism | 0.150 | 0.14 | 0.07 | 0.57 |
| M - membrane and cell wall structure and biogenesis | 0.140 | 0.09 | 0.05 | 0.36 |
| K - transcription | 0.140 | 0.09 | 0.05 | 0.36 |
| R - general functional prediction only | 0.140 | 0.09 | 0.04 | 0.36 |
| S - function unknown | 0.128 | 0.02 | 0.01 | 0.09 |
| G - carbohydrate metabolism and transport | 0.123 | -0.02 | -0.01 | -0.07 |
| U - intracellular trafficking and secretion | 0.122 | -0.02 | -0.01 | -0.09 |
| Q - biosynthesis, transport and catabolism of secondary metabolites | 0.112 | -0.10 | -0.05 | -0.40 |
| V - Defense | 0.106 | -0.16 | -0.08 | -0.62 |
| V(i) – Antibiotic/drug resistance | 0.135 | 0.06 | 0.03 | 0.25 |
| V(ii) – Anti-pathogen defense | 0.059 | -1.05 | -0.52 | -4.18 |
| Tr - transposon | 0.104 | -0.18 | -0.09 | -0.74 |
| Pl – conjugative plasmid | 0.079 | -0.53 | -0.27 | -2.12 |
| Ph – (pro)phage | 0.047 | -1.56 | -0.78 | -6.23 |

To shed light at the causes that make the defense genes slightly deleterious, we split the gene families in this category into two subcategories: (i) drug and/or antibiotic resistance and detoxification, (ii) restriction-modification, CRISPR-Cas and toxin-antitoxin. The median fitness effect substantially and significantly differs in sign and magnitude between both groups, with *s* = (3.1 × 10^−10^, 2.5 × 10^−9^) for genes involved in detoxification and drug resistance and *s* = (−4.2 × 10^−8^, −5.2 × 10^−9^) for genes involved in anti-parasite defense (Mann-Whitney test, *p* < 10^−7^). Thus, the drug resistance machinery is close to neutral whereas the anti-parasite defense systems are about as deleterious as plasmids and somewhat more so than transposons. Among the latter, toxin-antitoxins are the most deleterious, followed by CRISPR-Cas and restriction-modification, although the pairwise differences are only significant between toxin-antitoxins and restriction-modification (*s* = (−8.8 × 10^−8^, −1.1 × 10^−8^) and *s* = (−2.1 × 10^−9^, −2.6 × 10^−10^) respectively, Mann-Whitney test, *p* = 0.02).

### Long-term gene dynamics and bursts of transposon proliferation

The loss biases and selection coefficients in Table 1 describe the dynamics of genes in groups of closely related genomes, with evolutionary distances of approximately 0.01 to 0.1 fixed substitutions per base pair. To investigate whether the same values apply at larger phylogenetic scales, we pooled data from all ATGCs and compared the global abundances of genes from different categories with the long-term equilibrium abundances expected from the model (Fig.1B-C). In most categories, the observed copy number agrees with the predicted value, and the same holds for the fraction of genomes that harbor a given gene family.

Two notable exceptions are the genes involved in translation (category J) and the transposons. In the case of translation-related genes, the observed copy number is ~40% greater than expected (median observed 0.5, median expected 0.36, Wilcoxon test *p* < 10^−20^), and the fraction of genomes with at least one copy is ~80% greater than expected (median observed 0.48, median expected 0.27, Wilcoxon test *p* < 10^−20^). Such deviations reflect the inability of the model to reproduce a scenario in which selection acts to maintain a single member of most of the gene families in almost every genome, as is the case for translation. In the case of transposons, there is a dramatic excess of ~213% in the mean copy number (median observed 0.25, median expected 0.08, Wilcoxon test *p* < 10^−6^) but no significant deviation in the fraction of genomes that carry transposons. Such excess of copies apparently results from occasional proliferation bursts that offset the prevailing loss-biased dynamics. Indeed, ~12% of the lineage-specific families of transposons show evidence of recent expansions, as indicated by effective duplication/loss ratios greater than one, whereas the fraction of such families drops below 4% in other functional categories (Fig. 2A, orange bars). Analysis of the typical burst sizes also reveals differences between transposons, with a mean burst size close to 4, and the rest of genes, with mean burst sizes around 2 (Fig2A, gray line). Episodes of transposon proliferation are not evenly distributed among taxa, but rather concentrate in a few groups, such as *Sulfolobus*, *Xanthomonas*, *Francisella* and *Rickettsia* (Fig. 2B). The high prophage burst rate in *Xanthomonas* is due to the presence of a duplicated prophage related to P2-like viruses in *X. citri*.

**Figure 2:**
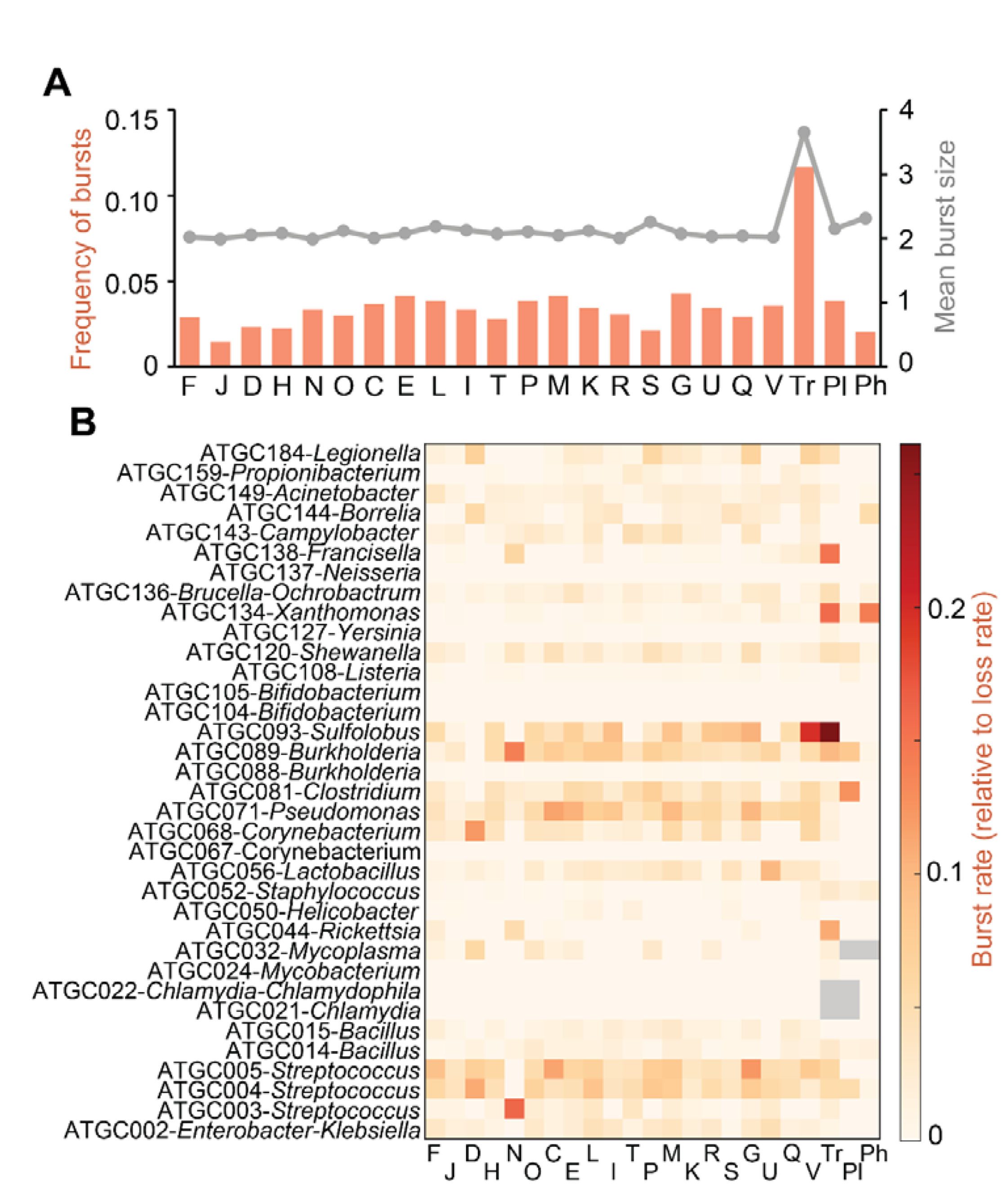
Frequency and distribution of proliferation bursts in different functional categories of genes. A: Orange (left axis), frequency of proliferation bursts, defined as the fraction of ATGC-COGs with effective duplication/loss ratio *d*/*l*_*e*_>1, split by functional category; gray (right axis), mean burst size for these ATGC-COGs. B: Burst rates in different ATGCs and functional categories, relative to the rate of gene loss. Designations of functional categories are the same as in Fig. 1 and Table 1.

To test whether the burst dynamics observed for transposons could explain the deviation in their global abundance, we analyzed a modified version of the model in which long phases of genome decay are punctuated by proliferative bursts of size *K*. Specifically, each decay phase was modeled as a duplication-transfer-loss process with selection, with initial condition *p*_*k*_ = 1, *p*_*k*≠*k*_ = 0. Bursts occur at exponentially distributed intervals with the rate *ϕ* (note that *T* − 1/*ϕ* is the characteristic interval between two consecutive bursts). When a burst occurs, the duplication-transfer-loss process is reset to its initial condition. In this model, the time-extended average for the mean copy number ≪k≫, becomes 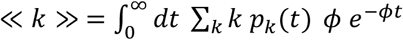. Using this expression it is possible to evaluate the expected mean copy number for any given value of the burst rate and the burst size (see SI text for details). In the case of transposons, the fraction of families with signs of recent expansions leads to the estimate *ϕ* = 0.04 (i.e one burst for every 25 losses, see Methods). For this burst rate, the modified model recovers the observed mean copy number if the burst size is set to *K* = 4.2, which is notably close to the value *K* = 3.9 estimated from the data.

### Relationships between genome size and gene duplication, horizontal transfer and loss rates

We further investigated the relationships between the genome size and the factors that determine gene abundances. For each set of related genomes, we estimated the intrinsic duplication/loss ratio (*d*/*l*) and the total HGT/loss ratio (*h*/*l*) for genes from ‘neutral’ categories and compared those to the mean genome size, quantified as the number of ORFs in the genome. As shown in Fig. 3, *d*/*l* is independent of the genome size, whereas *h*/*l* positively correlates with the genome size.

The same trends are confirmed by the analysis of ORFan abundances. Provided that the duplication rate is small compared to the loss rate, the number of ORFan families per genome constitutes a proxy for the ratio *h*/*l*. On the other hand, the fraction of ORFan families with more than one copy is a quantity that only depends on the ratio *d*/*l* (SI text). As in the case of neutral gene families, the study of ORFans reveals a strong positive correlation between genome size and *h*/*l*, but lack of significant correlation with *d*/*l*.

**Figure 3:**
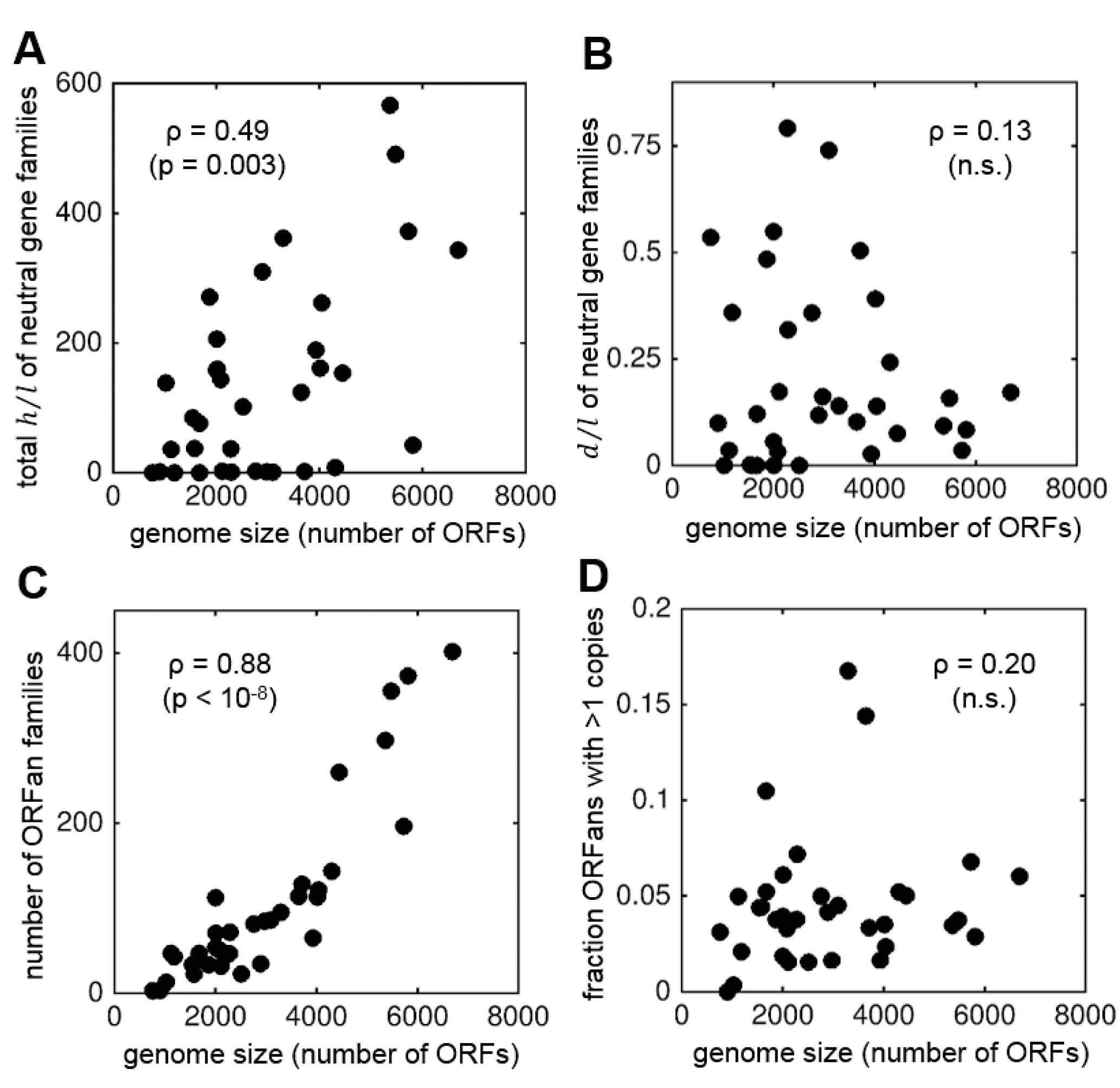
Correlations between the genome size and potentially relevant parameters of gene family dynamics and genome architecture. Each point represents an ATGC. A: Total HGT to loss ratio for genes from neutral categories. B: Duplication to loss ratio for genes from neutral categories (both duplication and loss rates are calculated per copy). C: Number of ORFan families per genome, which is an independent proxy for *h*/*l*. D: Fraction of ORFan families with more than one copy, which is proportional to *d*/*l*. In each panel, the Spearman’s *ρ* and significant *p*-values are shown; non-significant (n.s.) *p*-values are greater than 0.2.

Because in prokaryotes genome size positively correlates with the effective population size (*N*_*e*_) (14), we also explored the correlations between *N*_*e*_ and the ratios *h*/*l* and *d*/*l* (SI text). The same qualitative correlations were detected, i.e *h*/*l* positively correlates with *N*_*e*_, whereas *d*/*l* shows no correlation. However, the association between *h*/*l* and *N*_*e*_ becomes non-significant when genome size and *N*_*e*_ are jointly considered in an analysis of partial correlations. Therefore, it seems that the association between *N*_*e*_ and *h*/*l* is a by-product of the intrinsic correlation between effective population size and genome size.

### Disentangling environmental and intrinsic contributions to fitness

Because our estimates of the selection coefficients constitute ecological and temporal averages, a low selection coefficient might not only result from a genuine lack of adaptive value but, perhaps more likely, from the limited range of environmental conditions in which the given gene becomes useful. To disentangle the two scenarios, we compared the non-synonymous to synonymous nucleotide substitution ratios (*dN*/*dS*) for different gene categories. The expectation is that genes that perform an important function in a rare environment would be characterized by low average selection coefficients (frequent loss) combined with intense purifying selection at the sequence level (low *dN*/*dS*) in those genomes that harbor the gene. Gene sequence analysis shows that, in most cases, the *dN*/*dS* of a gene is primarily determined by the ATGC rather than by the functional category (SI Fig. S1). These observations are compatible with the results of a previous analysis indicating that the median *dN/dS* value is a robust ATGC-specific feature (15). Notable exceptions are transposons and prophages, which show a high *dN*/*dS* in most taxa.

After accounting for the ATGC-related variability, we found a significant negative correlation between the selection coefficient of a functional class and the *dN*/*dS* (Fig. 4, Spearman's *ρ* = −0.58, *p* = 0.004). Such a connection between the selection pressures on gene dynamics and sequence evolution is to be expected under the straightforward assumption that genes that are more important for organism survival are subject to stronger selection on the sequence level and has been observed previously (48). However, genes involved in metabolic processes, especially carbohydrate metabolism, have lower *dN*/*dS* values than predicted from the overall trend (Fig. 4), suggesting that the effective neutrality of such genes results from the heterogeneity of environmental conditions. Among the gene categories with low selection coefficients, the *dN*/*dS* values of transposons, prophages and gene families with poorly characterized functions are significantly greater than expected from the general trend, which is consistent with the notion that these genes provide little or no benefit to the cells that harbor them.

**Figure 4:**
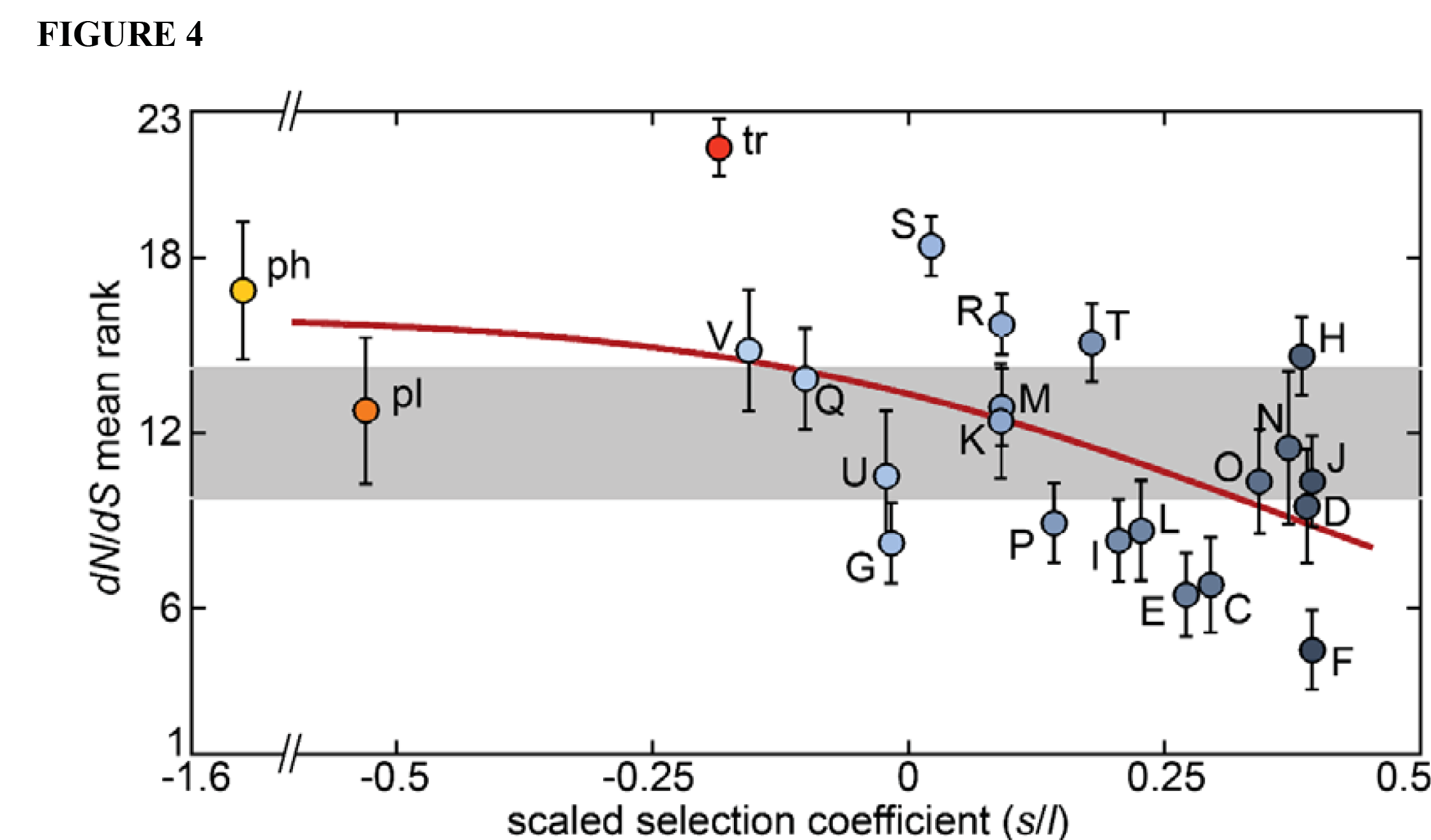
Comparison between the scaled selection coefficients (*s*/*l*) of different functional categories and their characteristic non-synonymous to synonymous mutation ratios (*dN*/*dS*). To account for ATGC-related variation, the *dN*/*dS* ratios for all categories within an ATGC were converted into ranks. Circles represent the mean ranks averaged across ATGCs, and error bars represent the standard error of the mean. Colors are the same as in Fig. 1. The horizontal grey band shows the theoretical 95% CI for the means of a null model where all categories have similar *dN*/*dS* (points above/below this interval indicate that the *dN/dS* of a category is significantly higher/lower than the expectation under the null model). The trend line (red) was obtained by fitting a monotonic spline curve to the data.

### Gene dynamics and microbial life styles

In an effort to clarify the biological underpinnings of the gene dynamics, we compared the effective duplication to loss ratios in microbes with three life styles: free-living, facultative host-associated and obligate intracellular parasite (Fig. 5). In the first two groups, *d*/*l*_*e*_ drops from essential functional categories to non-essential categories and genetic parasites, with significantly higher values in free-living microbes than in facultative host-associated bacteria. Obligate intracellular parasites have remarkably low *d*/*l*_*e*_ values, as could be expected from their strong genomic degeneration. Notably, genetic parasites and genes from the defense category show the highest *d*/*l*_*e*_ among the genes of intracellular parasites, although due to the small number of intracellular parasites in our dataset (only 3 ATGCs, with most genetic parasites restricted to the ATGC044 encompassing *Rickettsia*), this result must be taken with caution. We estimated the selection coefficients for free-living and facultative host-associated microbes, under the assumption that the intrinsic *d/l* is universally the same across the microbial diversity. The significant difference in *d*/*l*_*e*_ between the two lifestyles translates into consistently higher *s* values for most functional categories of genes in free-living microbes (SI Fig. S2). Thus, the beneficial effects of most genes appear to be significantly greater in free-living compared to facultative host-associated bacteria, and in both these categories of microbes, selection for gene retention is dramatically stronger than it is in obligate, intracellular parasites.

**Figure 5:**
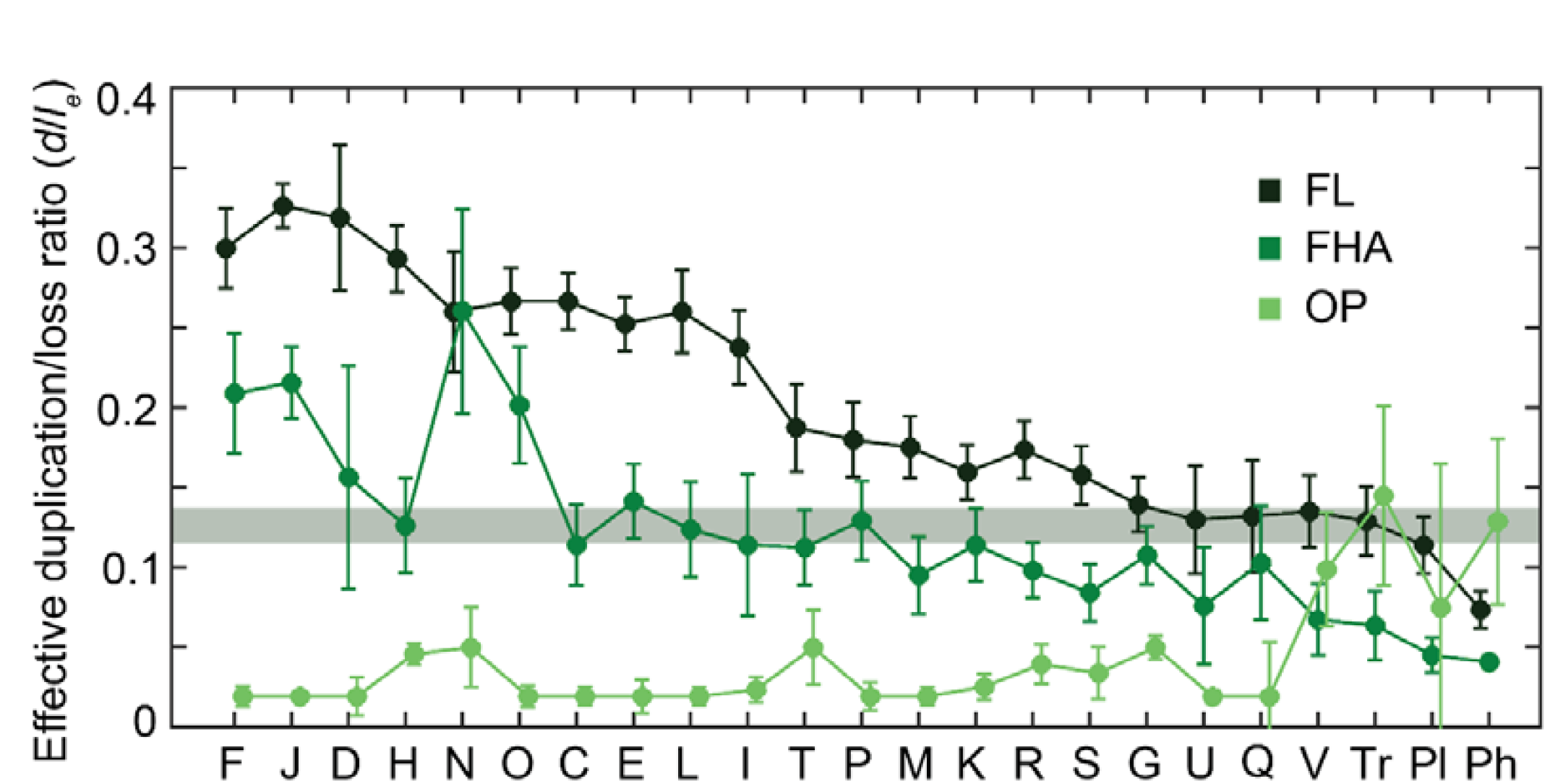
Effective duplication to loss ratio (*d*/*l*_*e*_) in free-living (FL), facultative host-associated (FHA) and obligate intracellular parasitic (OP) microbes. The designations of functional classes in the x-axis are the same as in Fig. 1 and Table 1. The shaded band indicates the 95% CI for the intrinsic *d/l* estimated from neutral categories and ORFans. Error bars denote the 95% CI for the median *d*/*l*_*e*_.

## Discussion

Multiple variants of the duplication-transfer-loss model and related multi-type branching processes have been widely used to study the evolution of gene copy numbers (24, 25, 28, 49), especially in the context of transposons and other genetic parasites (22, 23, 26, 27, 50). To make the models tractable, most studies make simplifying assumptions, such as stationary state, absence of duplication or lack of selection, and obtain the model parameters from the copy number distributions observed in large genomic datasets,relying on the assumption that model parameters are homogeneous across taxa. Here we derived an exact solution for the time-dependent duplication-transfer-loss model with additive selection and found that, in general, it is impossible to distinguish neutral and costly elements solely based on the copy number distributions. This is the case because the effects of selection and loss bias blend into a composite parameter that is equivalent to an effective loss bias in a neutral scenario. Using the solution of the complete model, we investigated the copy number dynamics of a large number of gene families in groups of related genomes, without the need to assume homogeneity of the HGT, duplication and loss rates across taxa (8). We then used the expression that relates the parameter values under selection with their neutral equivalents to estimate the selection coefficients for different classes of genes.

The results of this analysis rely on several assumptions. First, the duplication-transfer-loss model was solved in a regime of linear selection that is, assuming that the benefit or cost of a gene family linearly grows with the gene copy number. This choice of the cost function, that is arguably suitable for genetic parasites, might be violated by ensembles of genes involved in processes that require tight dosage balance among the respective proteins, such as the translation system (51). For such genes, the fitness benefit will be underestimated as the observed number of family members is lower than predicted by the model. Second, to calculate the intrinsic loss bias (*d*/*l*), we assumed that certain classes of genes are effectively neutral. In that regard, two independent approaches were explored: (i) using ORFans as the neutral class; (ii) inferring the neutral categories based on plausible dispensability and a low position in the effective loss bias ranking. Notably, nearly identical values were obtained through both approaches, indicating that our estimates are robust to the choice of the neutral reference group. Third, the model assumes that duplication and deletion rates, as well as selection coefficients, are constant in time. It has been proposed that recently duplicated genes are subject to significantly higher loss rates and lower selection coefficients than older paralogs (52, 53). Should that be the case, recently duplicated gene copies would be short-lived and their existence would not affect the generality of our results, provided that the duplication to loss ratio is understood as an effective parameter that accounts for the survival probability of a paralog beyond the initial phase. Finally, in order to convert the selection to loss ratios (*s*/*l*) to selection coefficients (*s*), we used two estimates of the loss rate *l*. A conservative estimate *l* = 5 × 10^−9^ was taken from the experimental study of medium to large deletions (in the range of 1 kb to 202 kb) in *Salmonella enterica* (37). Because small indels also contribute to the loss of genes via pseudogenization, we additionally considered a second, upper bound estimate, *l* = 4 × 10^−8^, which is the geometric mean of the indel rates collected from multiple mutation accumulation experiments (38-45) multiplied by an average target size of 1 kb per ORF.

Our estimates yielded a broad range of selection coefficients that reflects positive, near zero (neutral) or negative fitness contributions of the respective genes. Notably, the ranking of the gene categories by fitness contribution is closely similar to the ranking by evolutionary mobility (gene gain and loss rates) (8) such that genes with positive fitness contributions are the least mobile. In accordance with the intuitive expectation, gene families involved in essential functions, in particular nucleotide metabolism and translation, occupy the highest ranks in the list of genes maintained by selection (highest positive *s* values; Table 1). The middle of the range of selection coefficients is occupied by functional categories of genes that are beneficial, sometimes strongly so, for microbes under specific conditions but otherwise could be burdensome, such as carbohydrate metabolism and ion transport. This inference was supported by analysis of selection on the protein sequence level that is reflected in the *dN/dS* ratio. Overall, we observed the expected significant negative correlation between the selection coefficient estimated from gene dynamics and *dN/dS*, indicating that functionally important genes are, on average, subject to strong constraints on the sequence level. However, for genes involved in metabolic processes, in particular carbohydrate metabolism, the *dN/dS* values are lower than expected given their average selection coefficients, which is consistent with relatively strong sequence-level selection in the subsets of microbes that have these genes. In agreement with this interpretation, when the *s* values for these categories were estimated separately for free-living and host-associated microbes, they turned out to be slightly beneficial in the former but costly in the latter.

In contrast, genetic parasites that negatively contribute to the fitness of the cell are at the bottom of the list of *s* values (Table 1). Among those, prophages are the most costly class whereas plasmids and especially transposons evolve under regimes closer to neutrality. Prophages, plasmids and transposons differ substantially in the magnitude of the associated selection coefficients: selection is strong and effective against prophages (*N*_*e*_*s* ∼ −10), and moderate against transposons and plasmids (*N*_*e*_*s* ∼ −1). These differences are consistent with the differences in the lifestyles between these selfish elements whereby transposons and plasmids are relatively harmless to the host cell, apart from being an energetic burden, whereas prophages have the potential to kill the host upon lisogenization (20, 54). Accordingly, genetic parasites also differ in the relative importance that selection and deletions play in keeping them under control. Both selection and deletions contribute to the removal of prophages (the contribution of selection being ~1.6 times greater), whereas deletion is the main cause of plasmid and transposon loss (roughly twice as important as selection for plasmids and 5 times as important in the case of transposons). The demonstration that transposons are only weakly selected against and are lost primarily due to the intrinsic deletion bias is compatible with the wealth of degenerated insertion sequences found in many bacterial genomes (55-57). Conversely, deleterious elements, such as prophages, whose spread is limited by selection against high copy numbers, present fewer degenerated copies than lower cost elements, such as transposons.

One of the most interesting and, at least at first glance, unexpected observations made in the course of this work is that genes encoding components of anti-pathogen defense systems are on average deleterious, with an average cost similar to or even greater than the cost of plasmids (Table 1). In part, this is likely to be the case because some of the most abundant defense systems, such as toxin-antitoxins and restriction-modification modules, clearly display properties of selfish genetic elements and moreover, are addictive to host cells (58-61). Indeed, in agreement with the partially selfish character of such defense modules, we found that toxin-antitoxins are the most deleterious category of genetic elements in microbes, apart from prophages. More generally, the patchy distribution of defense systems in prokaryotic genomes, together with theoretical and experimental evidence, suggests that defense systems incur non-negligible fitness costs that are thought to stem primarily from autoimmunity and abrogation of HGT, and therefore, are rapidly eliminated when not needed (62-64).

Long-term transposon dynamics is well described by a model that combines long phases of decay, during which transposons behave as inactive genetic material, punctuated by small proliferation bursts that produce on average 4 new copies. Despite the simplicity of this model, it captures, at least qualitatively, the heterogeneity of transposition rates among transposon families (65) and environmental conditions (66, 67). Unlike large expansions, which are rare events typically associated with ecological transitions affecting the entire genome (68-71), small bursts occur frequently and affect a sizable fraction of transposon families. Some well-known instances of large transposon expansions become apparent in our analysis that identified taxa with unusually high burst rates, such as *Xanthomonas*, *Burkholderia* and *Francisella*, in accord with previous observations (70, 71). In most other taxa, transposon decay is the dominant process, which is the expected trend, given that transposition is tightly regulated and a large fraction of transposon copies are inactive (72, 73). The small fitness cost of transposons in the decay phase is also consistent with a non-proliferative scenario, where the fitness effect is reduced to the energetic cost of replication and expression (21). Due to the rapidity of bursts, our methodology cannot be used to assess the cost of a transposon during the burst phase. Because active transposons likely impose a larger burden on the host (74), variation in burst sizes is likely to reflect differences in the intensity of selection and the duration of proliferative episodes.

Apart from the transposons, the only notable case of burst-driven dynamics corresponds to genes from the defense category in *Sulfolobus*. A closer inspection of this group reveals multiple instances of duplications, gains and losses of CRISPR-Cas systems as also observed previously (75). In the case of prophages, the low burst rate is likely to reflect genuine lack of bursts or our inability to detect them due to the dominant, selection-driven fast decay dynamics. Indeed, given the fitness cost that we estimated for prophages, a burst of prophages would decay almost three times faster than a burst of transposons of similar size.

The effective size of microbial populations positively correlates with the genome size, which led to the hypothesis that the genome dynamics is dominated by selection acting to maintain slightly beneficial genes (14, 16). In the present analysis, when gene families from all functional categories are pooled, the median fitness contribution per gene is *N*_*e*_*s* ~0.1, which provides independent support for this weak selection-driven concept of microbial genome evolution. In that framework, the fact that genetic parasites are more abundant in large genomes, as reported previously (76-78) and confirmed by our data, seemingly raises a paradox: the same genomes where selection works more efficiently to maintain beneficial genes also harbor more parasites. A possible solution comes from our observation that the HGT to loss ratio (where the HGT rate is measured per genome and the loss rate is measured per gene) grows with the genome size. Such behavior, which had been already noted for transposons (23) and agrees with the recently derived genome-average scaling law (14), is likely to result, at least in part, from larger genomes providing more non-essential regions where a parasite can integrate without incurring major costs to the cell. Alternatively or additionally, the observed dependence could emerge if duplication and loss rates per gene decreased with genome size, while the HGT rate remains constant. Indeed, an inverse correlation between the genome size and the duplication and loss rates could be expected as long as mutation rates appear to have evolved to lower values in populations with larger *N*_*e*_ (41, 79).

Taken together, the results of this analysis reveal the relative contributions of selection and intrinsic deletion bias to the evolution of different classes of microbial genes and selfish genetic elements. Among other findings, we showed that the genome-averaged selection coefficients are low, and evolution is driven by strong selection only for a small set of essential genes. In addition, we detected substantial, systematic differences between the evolutionary regimes of bacteria with different lifestyles, with much stronger selection for gene retention in free-living microbes compared to parasites, especially obligate, intracellular ones. This difference appears to be fully biologically plausible in that diversification of the metabolic, transport and signaling capabilities is beneficial for free-living microbes but not for parasites that therefore follow the evolutionary route of genome degradation.

Counterintuitive as this might be, we show that anti-parasite defense systems are generally deleterious for microbes, roughly to the same extent as mobile elements. These results are compatible with the previously observed highly dynamic evolution of such systems that are kept by microbes either when they are essential to counteract aggressive parasites or due to their own selfish and addictive properties. These findings can be expected to foster further exploration of the interplay between genome size, effective population size, the rates of horizontal transfer, duplication and loss of genes and the dynamics of mobile elements in the evolution of prokaryotic populations and, eventually, the entire microbial biosphere.

## Methods

### Gene copy number dynamics

Let *n*_*k*_(*t*) be the number of genomes that carry *k* copies of the gene of interest at time *t*. We define the generating function 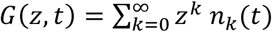. In terms of the generating function, eq. (1) becomes 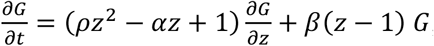, where 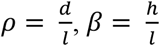, and 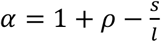. This equation can be solved for any initial condition by applying the method of characteristics (SI text). The generating function for the copy number distribution *p*_*k*_(*t*) is then obtained as *H*(*z*, *t*) = *G*(*z*, *t*)/*G*(1, *t*). The explicit values of *p*_*k*_(*t*) are recovered as the coefficients of the series expansion of *H*(*z*, *t*) with respect to *z*.

### Estimation of the effective ratios *d*/*l*_*e*_ and *h*/*l*_*e*_ from genomic data

Genomic data were obtained from an updated version of the ATCG database that clusters genomes from bacteria and archaea into closely related groups (3). We analyzed 35 of the largest ATGCs (34 bacterial and one archaeal group) that included 10 or more genomes each. For each of those ATGCs, Clusters of Orthologous Genes shared among genomes of the same ATGC (ATGC-COGs) were identified (33, 80) and rooted species trees were generated as described previously (8).

The effective duplication/loss ratio (*d*/*l*_*e*_) and transfer/loss ratio (*h*/*l*_*e*_) for each ATGC-COG were estimated with the software COUNT (24), which optimizes the parameters of a duplication-transfer-loss model analogous to the model described above under the assumption of neutrality (81). The output of the program was post-processed to obtain ATGC-COG-specific rates as described in (20). ATGC-COGs were assigned to families based on their COG and pfam annotations. COG and pfam annotations were also used to classify families into functional categories. At the family level, the representative ratios *d*/*l*_*e*_ and *h*/*l*_*e*_ of a family were obtained, respectively, as the median *d*/*l*_*e*_ and the sum of *h*/*l*_*e*_ among its constituent ATGC-COGs. The mean copy number of a family was calculated as the average, across all ATGCs, of the ATGC-specific mean abundances (ATGC-COGs belonging to the same family in the same ATGC were pooled to obtain the ATGC-specific mean abundance, whereas the ATGC-specific mean for absent families was set to zero). The fraction of genomes that contain a family was calculated in a similar manner. This approach minimizes the bias associated to non-uniform ATGC sizes. To minimize inference artifacts associated to small families, only those families encompassing at least 5 ATGC-COGs from at least 3 ATGCs were considered for further analyses.

### Estimation of the intrinsic duplication/loss ratio

Two approaches were used to estimate the intrinsic duplication/loss ratio *d*/*l*. In the first approach, putative neutral families from categories R, S, G, U, Q, and V were pooled and the median *d*/*l*_*e*_ was chosen to serve as the estimate of *d*/*l*. The 95% confidence interval was calculated with the formula *median* 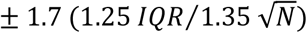, where *IQR* is the interquartile range and *N* is the number of families (82). In the second approach, the copy numbers of ATGC-COGs that are specific to one single genome were used to infer the ratio *d*/*l* under the assumption that such genes are of recent acquisition and effectively neutral. To that end we used the solution of the duplication-transfer-loss model to derive a maximum likelihood estimate of *d*/*l* given a list of single-genome ATGC-COGs, their copy numbers, and the time since the last branching event in the genome tree (in units of loss events, as provided by COUNT). Explicit formulas and their derivation are discussed in the SI text. Likelihood maximization was carried out using the Nelder-Mead simplex method as implemented in MATLAB R2016b. The 95% confidence interval was determined by the values of *d*/*l* whose log-likelihood was 1.92 units smaller than the maximum log-likelihood (83).

### Burst frequency, rate and size

The frequency of bursts was calculated as the fraction of ATGC-COGs in which *d*/*l*_*e*_ > 1. The burst rate *ϕ* was estimated by maximum likelihood, assuming that bursts occur randomly at exponentially distributed intervals, such that the probability of observing a burst in a tree of phylogenetic depth *t* is equal to 1 − *e*^−*ϕt*^. Accordingly, the log-likelihood of observing *n_atgc_* bursts in an ATGC with *N_atgc_* ATGC-COGs 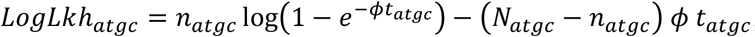, where *t*_*atgc*_ is the depth of the ATGC tree in units of loss events (SI text). The global log-likelihood is the sum of the contributions from all ATGCs. As a proxy for the burst size we used the maximum copy number observed in each ATGC-COG. For each category, the characteristic burst size was calculated as the quotient between the mean burst size in ATGC-COGs with *d*/*l*_*e*_ > 1 and the baseline defined by the mean of the maxima in the rest of ATGC-COGs.

### Estimation of the characteristic *dN*/*dS* ratios

The *dN*/*dS* of every ATGC-COG was calculated as follows. Starting from the multiple sequence alignment, the program codeml from the PAML package (84) was used to obtain the *dN*/*dS* for each pair of sequences in the ATGC-COG. The conditions 0.01 < *dN* < 3 and 0.01 < *dS* < 3 were used to select informative gene pairs. The representative *dN*/*dS* for the ATGC-COG was obtained as the median *dN*/*dS* among the informative pairs. In the next step, ATGC-COGs from the same ATGC that belong to the same functional category were pooled and the median of their *dN*/*dS* was taken as the representative *dN*/*dS*. To account for ATGC-related effects, the *dN*/*dS* values of all categories within an ATGC were converted into ranks. The null hypothesis that all categories are equal in terms of their *dN*/*dS* was rejected by a Skillings-Mack test (*t* = 939.7, d.f. = 22, *p* < 10^−20^). To identify which categories significantly deviate from the null hypothesis, the mean rank of each category was compared to the theoretical 95% CI for the mean of 35 samples taken from a discrete uniform distribution in the interval from 1 to 23.

